# Structural insights as to how iron-loaded siderophores are imported into *Mycobacterium tuberculosis* by IrtAB

**DOI:** 10.1101/2022.01.11.475891

**Authors:** Shan Sun, Yan Gao, Xiaolin Yang, Xiuna Yang, Tianyu Hu, Jingxi Liang, Zhiqi Xiong, Pengxuan Ren, Fang Bai, Luke W. Guddat, Haitao Yang, Zihe Rao, Bing Zhang

**Affiliations:** Shanghai Institute for Advanced Immunochemical Studies and School of Life Science and Technology, ShanghaiTech University, Shanghai, 201210, China; State Key Laboratory of Medicinal Chemical Biology, Nankai University, Tianjin, 300353, China; Laboratory of Structural Biology, Tsinghua University, Beijing, 100084, China; National Laboratory of Biomacromolecules, CAS Center for Excellence in Biomacromolecules, Institute of Biophysics, Chinese Academy of Sciences, Beijing, 100101, China; School of Chemistry and Molecular Biosciences, The University of Queensland, Brisbane, QLD 4072, Australia; CAS Center for Excellence in Molecular Cell Science, Shanghai Institute of Biochemistry and Cell Biology, Chinese Academy of Sciences, Shanghai, 200031, China; University of Chinese Academy of Sciences, Beijing, 100101, China; Shanghai Clinical Research and Trial Center, Shanghai, 201210, China

**Keywords:** ABC exporter-like importer, Iron-loaded siderophore, IrtAB, Asymmetric conformational change, *Mycobacterium tuberculosis*

## Abstract

The ATP-binding cassette (ABC) transporter, IrtAB, plays a vital role in the replication and viability of *Mycobacterium tuberculosis* (*Mtb*), where its function is to import iron-loaded siderophores. Unusually, it adopts the typical fold of canonical ABC exporters. Here, we report the structure of unliganded *Mtb* IrtAB and its structure in complex with ATP, ADP, and an ATP analogue (AMP-PNP) at resolutions from 2.9 to 3.5 Å. In the ATP-bound state, the two nucleotide-binding domains (NBDs) form a “head-to-tail” dimer, but IrtAB has an unexpectedly occluded conformation, with the inner core forming a large hydrophilic cavity of about 4600 Å^3^. Comparison of the structure of the transporter in inward-facing and occluded conformations reveals that the NBD and the intracellular helical region of transmembrane domain (TMD) have an asymmetric allosteric mechanism when ATP binding/hydrolysis such that the one exhibits rigid-body rotation and the other moves in a concerted response as a rigid body. This study provides a molecular basis for the ATP-driven conformational changes that occur in IrtAB and an explanation as to how iron-loaded siderophores are imported into *Mtb* by IrtAB.

## Introduction

Iron, as an enzyme cofactor, is an essential nutrient required in almost all kingdoms of life including the human pathogen, *Mycobacterium tuberculosis* (*Mtb*) (De Voss et al., 2000; Rodriguez and Smith, 2006; Ryndak et al., 2010). However, for *Mtb* the amount of free iron is severely limited by resources in the human host (Begg, 2019; Skaar, 2010), since it is mainly stored in metal binding proteins such as ferritin (Arosio et al., 2017), lactoferrin (Brock, 2012) transferrin (de Jong et al., 1990) and intracellular heme (Paoli et al., 2002). In order to scavenge iron from the human host and in-turn to promote infection and pathogenicity (De Voss et al., 2000; Wells et al., 2013), *Mtb* possesses two classes of high-affinity iron siderophores (Gobin et al., 1995; Snow and White, 1969). These are lipid-bound mycobactin (MBT) and a secreted water soluble variant, carboxymycobactin (cMBT) (Gobin et al., 1995; Snow and White, 1969).

The heterodimeric complex IrtAB has been identified as being responsible for the import of the two iron-loaded siderophores (Fe^3+^-MBT and Fe^3+^-cMBT) from the periplasmic space into the cytoplasm of *Mtb* and other mycobacteria (Arnold et al., 2020; Rodriguez and Smith, 2006). This protein complex is vital for the replication and viability of *Mtb* in macrophages, which reside in the alveoalar (Rodriguez and Smith, 2006; Ryndak et al., 2010). IrtAB belongs to the adenosine 5′-triphosphate (ATP)– binding cassette (ABC) transporter superfamily (Braibant et al., 2000). Its function of importing siderophores is driven by the binding and hydrolysis of ATP (Arnold et al., 2020; Rodriguez and Smith, 2006). Like other ABC transporters, IrtAB contains two transmembrane domains (TMDs) and two nucleotide-binding domains (NBDs), with a single TMD and NBD present in each of IrtA and IrtB (Braibant et al., 2000). A unique feature of IrtAB is that IrtA contains a N-terminal siderophore interaction domain (SID) that controls the reduction of Fe^3+^-MBT by an electron donor, flavin-adenine dinucleotide (FAD), and this domain is essential for the uptake of Fe^3+^-MBT (Arnold et al., 2020; Ryndak et al., 2010). In contrast, the SID of IrtA has little effect on Fe^3+^-cMBT acquisition (Arnold et al., 2020). Interestingly, although IrtAB functions as an ABC importer, it adopts the canonical fold of type IV ABC exporters (Arnold et al., 2020), which are normally responsible for the efflux of intracellular substrates such as the multidrug resistance protein, Sav1866 (Dawson and Locher, 2006). In addition, IrtAB does not require a substrate binding protein (SBP) to capture specific substrates in the periplasmic space (Arnold et al., 2020), whereas canonical ABC importers such as maltose transporter, MBP-MalFGK_2_ (Oldham et al., 2007), and trehalose transporter, LpqY-SugABC (Liu et al., 2020), require this assistance. Therefore, IrtAB can be classified as belonging to a new subfamily of ABC importers with a distinct transport mechanism.

Based on multiple crystal and cryo-electron microscopy (cryo-EM) structures, the transport mechanism of canonical ABC exporters and importers is well understood (Chen, 2013; Hofmann et al., 2019; Locher, 2016; Mi et al., 2017; Rice et al., 2014). Despite the fact that structures for three transporters, YbtPQ (Wang et al., 2020), Rv1819c (Rempel et al., 2020) and ABCD4 (Xu et al., 2019) have been determined, the precise details as to how these ABC exporter-like importers, especially importers with a type IV exporter fold, operate have remained elusive. To date, there is also limited information to explain how IrtAB mediates the import of iron-loaded siderophores across the membrane in *Mtb*. This is highlighted by the fact that, amongst all mycobacteria, only the crystal structure of SID truncated IrtAB and the low resolution (6.9 Å) cryo-EM structure of full-length IrtAB (both all from *Mycobacterium thermoresistibile*) have been determined (Arnold et al., 2020). Therefore, in order to fully understand the complete functionality of IrtAB more structures in differently liganded states are required.

Herein, we have used cryo-EM to determine four structures of *Mtb* IrtAB in different states (Supplementary Table 1). These structures provide valuable insights into the molecular basis for the acquisition of iron-bound siderophores by pathogens.

## Results

### Biochemical characterization

Full-length *Mtb* IrtAB was overexpressed in *Mycobacterium smegmatis* (*M. smegmatis*) (strain mc^2^155), purified in lauryl maltose neopentyl glycol (LMNG) and exchanged into digitonin detergent (Supplementary Figure 1A). Two-dimensional (2D) class averages of full-length IrtAB cryo-EM particle images showed clear structural features with the SID of IrtA clearly visible in some 2D class orientations (Figure 1A and Supplementary Figure 2). However, there was no SID observed in the final three-dimensional (3D) reconstruction of this unliganded structure of the protein (Figure 1B and Supplementary Figure 2). It is suspected that this may be related to the long flexible loop at the join between the SID and the first elbow helix of the transmembrane (TM) region of IrtA (Supplementary Figure 3A). This flexible link could cause the SID of IrtA to swing relative to the TM region. Considering that the SID of IrtAB is not essential for transport of Fe^3+^-cMBT across the membrane in mycobacteria (Arnold et al., 2020), and the random swinging of SID could influence structural studies, we also expressed and purified SID truncated IrtAB (IrtAB_ΔSID_) (Supplementary Figure 1C). The ATPase activities of the full-length IrtAB and SID deletion construct (IrtAB_ΔSID_) were also measured and shown to be similar (Supplementary Figure 1I), which is consistent with the SID of IrtAB having little effect on Fe^3+^-cMBT uptake (Arnold et al., 2020).

**Figure 1.**
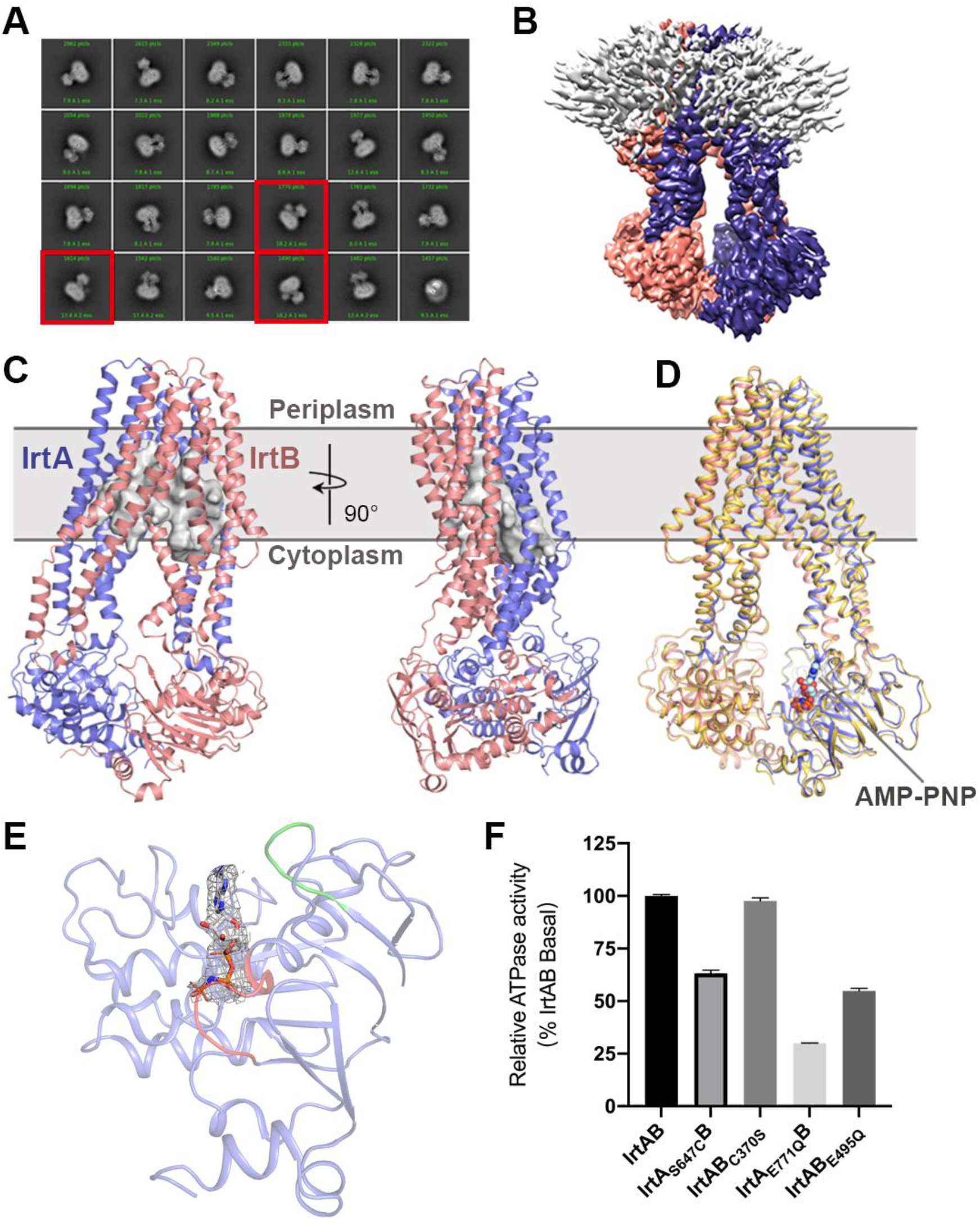
Cryo-EM structures of IrtAB from *M. tuberculosis*. (**A**) Representative 2D classification averages showing full-length IrtAB in different orientations. The SID of IrtA is observed in some views indicated by the red boxes. (**B**) Cryo-EM map of full-length IrtAB. IrtA is in slate, IrtB is in salmon, the detergent micelle is in gray. (**C**) Cartoon representation of inward-facing IrtAB viewed from the plane of the membrane. The membrane is shown in gray. IrtA and IrtB are in slate and salmon, respectively. The cavity of IrtAB is shown as a gray surface. (**D**) Superposition of ATP-free (yellow) and AMP-PNP-bound (IrtA, slate; IrtB, salmon) structures. The ATP analogue AMP-PNP is drawn as spheres and binds to the NBD of IrtA. (**E**) Cartoon representation of NBD of IrtA binding AMP-PNP. The Walker A and A-loop are colored red and green, respectively. The density map of AMP-PNP is shown as gray mesh and contoured at 9 σ. (**F**) The ATPase activities of four IrtAB variants were normalized relative to that of the wild-type (WT) IrtAB. Error bars represent mean ± SD based on three independent measurements.

### The cryo-EM structures of *Mtb* IrtAB, inward-facing states

The cryo-EM structure of full-length IrtAB without a bound nucleotide resembles the canonical fold of type IV ABC exporters (Mi et al., 2017; Perez et al., 2015), but its TM region has a partially collapsed inward-facing configuration (Figure 1C). The overall fold of *Mtb* IrtAB is similar to that of the crystal structure of the SID truncated counterpart from *Mycobacterium thermoresistibile* (Arnold et al., 2020) (RMSD of 2.38 Å after superimposition of 1021 Cα atoms) (Supplementary Figure 3B). However, there are several differences between these two structures, which may be, in part, due to the different methods used for structure determination. For the IrtA subunit, the differences are mainly located in the TM region, where the transmembrane helices have different degrees of shift at the cytoplasmic and periplasmic sides (Supplementary Figure 3C). It is noteworthy that TMH4 (flanking helix) and TMH5 have the largest conformational changes. Moreover, TMH3 also shows an increased displacement at the cytoplasmic side. By contrast, the TM region and NBD of IrtB are both different in these two structures (Supplementary Figure 3D). The changes to the TM region in IrtB are similar to that observed in IrtA. Moreover, the flanking helix (TMH4) is broken at the cytoplasmic side in our cryo-EM structure. The NBDs of IrtB show a significant translational shift, which could be related to the binding of the antibody at this position in the crystal structure.

We next determined the structure of IrtAB_ΔSID_ at 2.9 Å resolution in the presence of 1mM AMP-PNP, which is an inert ATP analogue, adenosine 5′-(β,γ-imido)triphosphate (Figure 1D and Supplementary Figure 4). Surprisingly, IrtAB still adopts an inward-facing conformation similar to the nucleotide-free full-length structure (Figure 1D). The density for AMP-PNP is observed near the A loop and Walker A motif of the NDB only in IrtA (Figure 1D, E). It is not present in IrtB. Superimposition of full-length IrtAB and this structure (IrtAB_ΔSID_-AMP-PNP bound) gives an RMSD of 0.85 Å for 1074 Cα atoms aligned, which suggests the SID of IrtAB has little effect on the structures of the TM regions and NBDs for this transporter. The structure of the complex further shows that there is asymmetric binding of the nucleotides in IrtAB, and implies that the NBDs of IrtA and IrtB have different binding affinities for nucleotides such as ATP. Such a situation has also been observed in the heterodimeric ABC exporter, TM287/288 (Hohl et al., 2012). In that case, the NBDs contain one degenerate site and one consensus site (Hohl et al., 2012; Procko et al., 2009). It has also been shown that the variations between the second and third glycine residues of the Walker A motifs in NBD domains can strongly affect the binding/hydrolysis of ATP in the multidrug resistance-associated protein (MRP1) (Yang et al., 2003). Sequence analysis showed that Ser647 of the Walker A motif in IrtA is replaced by Cys370 at the corresponding position in IrtB (Supplementary Figure 5), which implies that IrtA and IrtB may have different roles in ATP binding/hydrolysis as is observed in MRP1. To test this hypothesis, we interchanged these two residues in the two NBDs of IrtAB and measured the ATPase activities of these two mutants (IrtA_S647C_B and IrtAB_C370S_). The results showed that the S647C mutation greatly decreased ATP hydrolysis in IrtAB compared to the wild-type protein, whereas the C370S mutation in IrtB had almost no effect on ATPase activity (Figure 1F). In addition, the inactivation of ATPase activity in IrtA has a greater impact on the complex compared to IrtB (Figure 1F). Taken together, these results suggest the NBDs of IrtA and IrtB have different roles in ATP binding/hydrolysis and possibly have different functions during the uptake of iron-loaded siderophores.

### Structure of an IrtAB mutant in complex with ATP has an occluded state

To trap IrtAB in its ATP-bound state, the catalytic glutamate residues (Glu771 in IrtA and Glu495 in IrtB) were replaced by glutamine (IrtAB (E-Q)). We firstly tried to determine the structure of full-length IrtAB (E-Q) in complex with ATP, but this sample was unstable under freezing conditions. By contrast, the SID truncated IrtAB (E-Q) (IrtAB_ΔSID_ (E-Q)) incubated with ATP was suitable for cryo-EM and could be resolved at 3.1 Å resolution (Figure 2A and Supplementary Figure 6, 7A and Supplementary Table 1). Strong densities for ATP-Mg^2+^ bound to IrtA and IrtB are visible, resulting in an overall conformation that is substantially different from that of the ATP-free form. Due to the binding of ATP-Mg^2+^, the conformations of IrtA and IrtB are similar with an RMSD of 2.59 Å after the superimposition of 309 Cα atoms (Supplementary Figure 7B).

**Figure 2.**
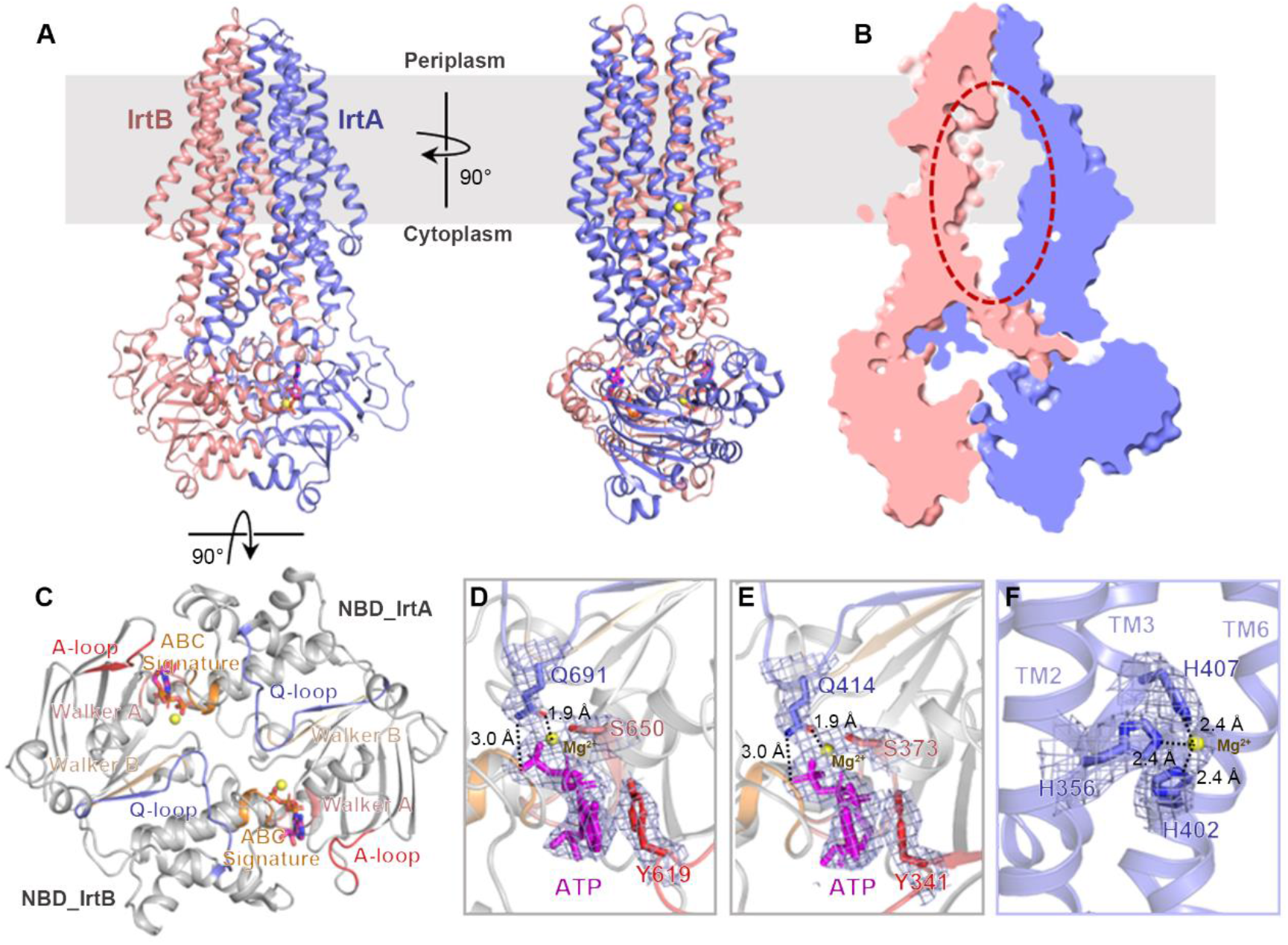
Cryo-EM structure of IrtAB_ΔSID_ (E-Q) in complex with ATP. (**A**) Two orthogonal views of the ATP-bound IrtAB_ΔSID_ (E-Q). ATP is represented as a stick model colored magenta, and Mg^2+^ ions are shown as yellow spheres. (**B**) Slice-through representation of the internal cavity of IrtAB. The hydrophilic cavity is indicated by a dashed red ellipse. (**C**) View of the NBD dimer from the periplasm. The highly conserved sequence motifs are also indicated. (**D**) Molecular interactions at the IrtA ATPase site, together with the density map (mesh) for ATP-Mg^2+^ and nearby residues. Conserved key residues are drawn as stick models. Interactions between glutamine in the Q loop and ATP or Mg^2+^ are indicated by dashed lines. (**E**) Molecular interactions at the IrtB ATPase site, together with the density map (mesh) for ATP-Mg^2+^ and nearby amino acids. Conserved key residues are drawn as stick models. Interactions between glutamine in the Q loop and ATP or Mg^2+^ are indicated by dashed lines. (**F**) Close-up view of the metal ion binding site in the TM region of IrtA. The density map for Mg^2+^ and residues that ligand to this metal is shown in mesh. Interactions between histidine and Mg^2+^ are indicated by dashed lines.

A comparison of before and after ATP binding shows that the TM helices of IrtAB do not split into two wings when ATP binds (Figure 2A, 3A). This is in contrast to the human lysosomal cobalamin transporter, ABCD4 (Xu et al., 2019), a trans-acting exporter responsible for efflux of cobalamin from the lysosome to the cytosol, where there is a lysosome-open conformation structure upon ATP binding. In IrtAB, the TM helices pack closely together to form an occluded cavity with an approximate volume of ∼4600 Å^3^ (Figure 2B), which spans the entire thickness of the membrane and extends to the intracellular part of IrtAB. Therefore, the structure determined here represents an intermediate state between the inward- and outward-facing conformation, similar to that of the ABC exporter McjD in complex with AMP-PNP (Choudhury et al., 2014). Surprisingly, density for a metal ion liganded to His356 (TMH2), His402 (TMH3) and His407 (TMH3) located inside the cavity of IrtA is observed (Figure 2F). These three residues are highly conserved across all mycobacteria. Considering 10 mM Mg^2+^ was added to the purification buffer, it is most likely that this is the identity of the metal ion, but its precise determination needs further studies. Since the metal ion is located near the cavity in IrtA (Supplementary Figure 8C), we suggest it may be involved in the stabilization of the substrate binding cavity during the process of transport.

At the cytoplasmic side, the two NBDs of IrtA and IrtB form a “head-to-tail” dimer, characteristic of ABC transporters (Figure 2A, C). Two ATP molecules are bound at the dimer interface, each of which is stabilized by an interaction between a Walker A motif in one NBD (IrtA or IrtB) and an ABC signature motif in the other NBD (IrtB or IrtA) (Figure 2C). Two magnesium ions are also observed in this structure, which ligand between the β- and γ-phosphates of ATP, a serine in the Walker A motif and the Q-loop glutamine, near the ABC signature motif (Figure 2D, E). The structures of the two NBDs of IrtAB are similar (RMSD of 1.1 Å after superimposition of 194 Cα atoms), with the differences mainly located in the α-helical subdomains (Supplementary Figure 7C), which is unique to ABC transporters and generally is structurally more diverse than the other subdomains (ter Beek et al., 2014). The contacts of two ATP molecules with nearby residues are very similar in IrtAB (Supplementary Figure 7D, E). Upon ATP binding, the density for the C-terminal extension of the NBD of IrtB is not observed, thus this region may have undergone conformational changes, suggesting it is directly related to ATP binding.

### Conformational changes upon ATP binding

Upon ATP binding, IrtA and IrtB rotate toward the center of the IrtAB complex, but these conformational changes are asymmetric with the largest displacement observed at the C-terminal ends of the NBDs of IrtA and IrtB, being ∼10 Å and ∼4 Å, respectively (Figure 3A), which is consistent with the conformational changes to the coupling helix (IH2) in IrtB and IrtA (Figure 3B, C). The conformational changes of IrtA and IrtB are mainly located in TMHs in the membrane inner leaflet and the NBDs in the cytoplasm, while IrtA has larger changes in comparison with that of IrtB (Figure 3A). For IrtA, TM helices 2-6 undergo rigid body conformational changes and TM helices 2-3 and TMH6 move towards each other with TM helices 4-5 blocking the lateral opening at the side of IrtA (Figure 3B). TM helices 4-5 of IrtB undergo a similar conformational change to that of IrtA in order to close the lateral opening at the IrtB side (Figure 3C). In addition to these global movements, at the binding site of the metal ion in the TM region of IrtA, the three histidine residues (His356, His402 and His407) undergo local conformational changes to optimally coordinate to the metal ion (Figure 3D). Notably, TMH5 of IrtB undergoes structural rearrangements before and after ATP binding at the cytoplasmic side (Figure 3E), which is a unique feature not reported previously in ABC transporters. It is commonly found that TMH4 (flanking helix) adjacent to TMH5 in ABC exporters undergoes structural rearrangements in different transition states in order to control the lateral opening in the membrane inner leaflet (Jin et al., 2012; Kim and Chen, 2018; Kodan et al., 2014). Here, the conformational changes of TMH5 in IrtB may contribute to stabilizing the interaction interface between NBD of IrtA and TMD (NBD_IrtA_-IH2_IrtB_), that is, Arg216 in TM5 of IrtB interacts with Asp680 of the RecA-like subdomain in NBD of IrtA by forming a salt bridge (Figure 4A).

**Figure 3.**
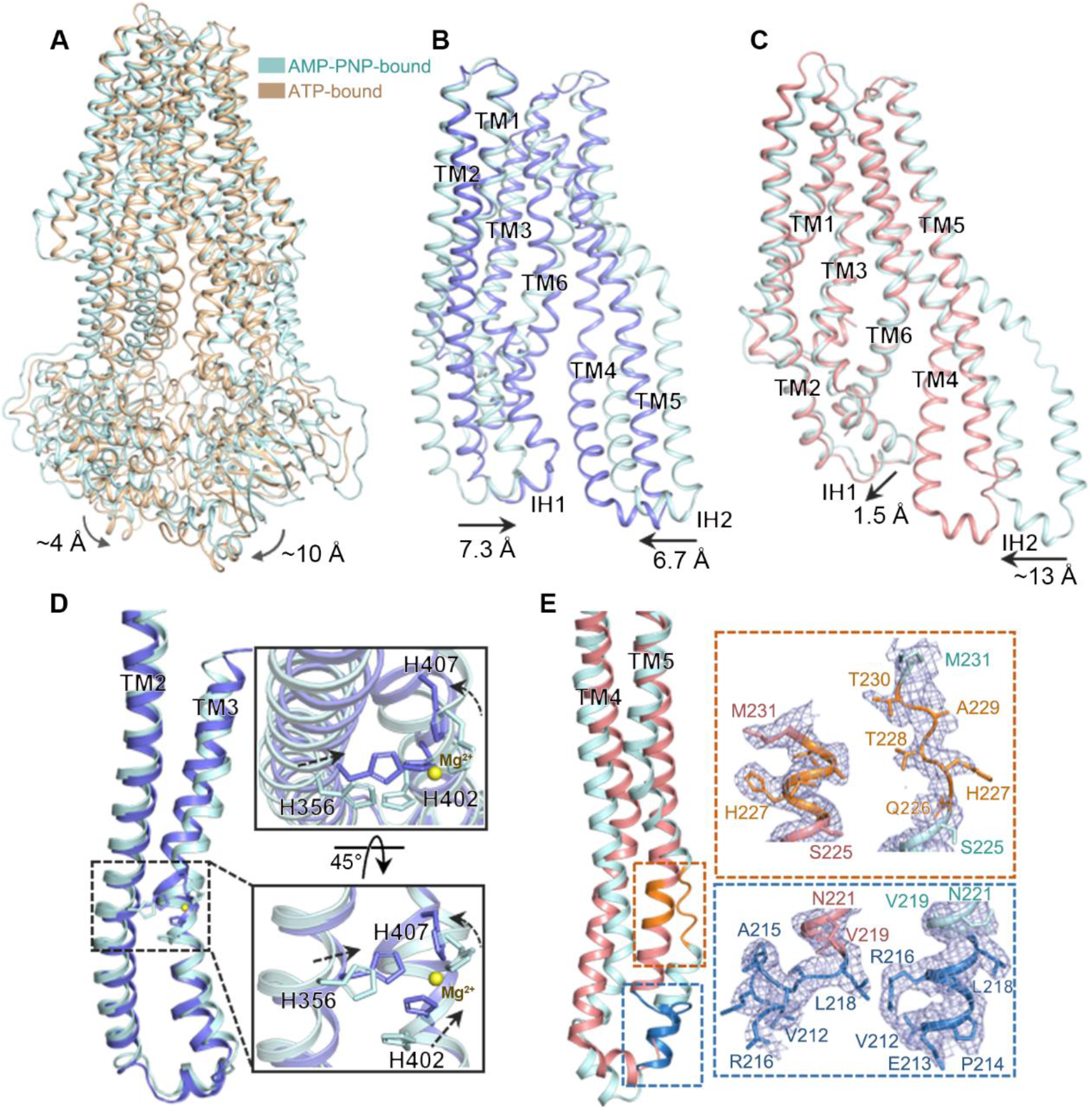
Conformational changes to IrtAB upon ATP binding. (**A**) Superposition of ATP-bound (gold) and AMP-PNP-bound (palecyan) structures. (**B**) Superposition of the TM region of IrtA in ATP-bound state (slate) and AMP-PNP-bound state (palecyan). The shift of the intracellular helices (IH) is indicated. (**C**) Superposition of the TM region of IrtB in ATP-bound state (salmon) and AMP-PNP-bound state (palecyan). The shift of the intracellular helices (IH) is indicated. (**D**) Close-up view of the conformational changes of metal ion binding site in TM region of IrtA upon ATP binding. (**E**) The structural rearrangements of TMH5 in IrtB upon ATP binding.

**Figure 4.**
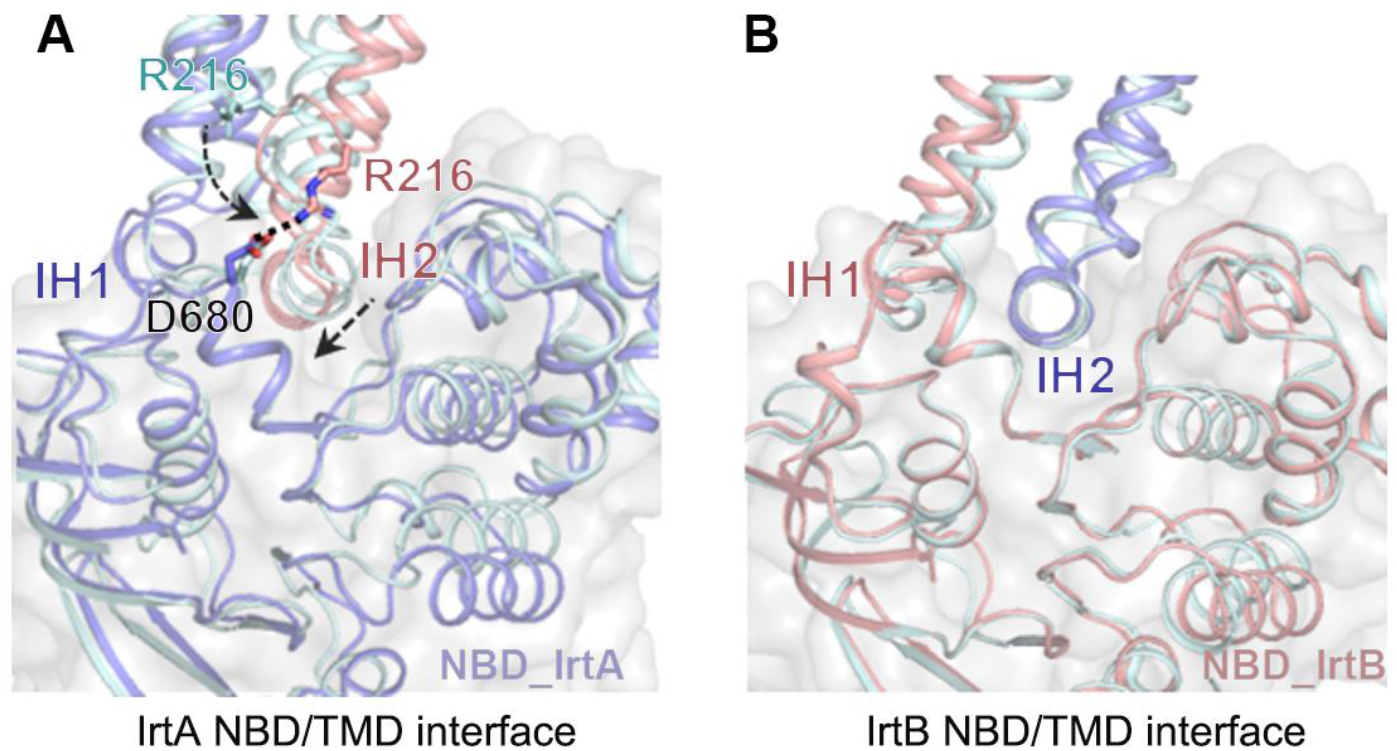
Asymmetric conformational changes at the TMD/NBD interface. (**A** and **B**) Cartoon representation of NBD/TMD interface in IrtA (A) and IrtB (B) before and after ATP binding. Structures of the inward-facing IrtAB (AMP-PNP-bound, palecyan) and the occluded IrtAB (ATP-bound) are superposed with respect to the NBDs. The surface clefts where the intracellular helices are docked into, are outlined in gray. The shift of IH2 in IrtB and the conformational change of Arg216 in TM5 of IrtB are also indicated. Interaction between Arg216 in IrtB and Asp680 in IrtA is indicated by the dashed line.

The NBD/TMD interface of the ABC transporter is crucial in coupling the conformational changes during ATP binding and hydrolysis (Kim and Chen, 2018). Multiple crystal structures of the maltose importer, MBP-MalFGK_2_, indicate that the coupling helix rotates relative to the NBDs during the transport cycle (Khare et al., 2009), while studies of P-glycoprotein (P-gp) showed that the NBD and the intracellular helical region of the TMD move as one concerted rigid body in the transport cycle (Esser et al., 2017; Kim and Chen, 2018). In contrast, the two equivalent interfaces in IrtAB (NBD_IrtA_-IH2_IrtB_ and NBD_IrtB_-IH2_IrtA_) show different features during the switch from the inward-facing to occluded conformation (Figure 4), that is, the NBD_IrtA_-IH2_IrtB_ interface is similar to that of the maltose importer, where the IH2_IrtB_ rotates relative to the NBD_IrtA_, whereas the NBD_IrtB_-IH2_IrtA_ interface is largely unchanged, similar to that of P-glycoprotein, where NBD_IrtB_-IH2_IrtA_ moves as one concerted rigid body. Therefore, in the transport cycle, the NBD and the intracellular helical region of TMD in IrtAB exhibit an asymmetric allosteric mechanism that is novel amongst the ABC transporter family.

## Discussion

Although the ABC transporter IrtAB plays a vital in the replication and viability of *Mtb* in human macrophages, we are at an early stage in understanding its molecular mechanism. In addition to determination of the structures of unliganded IrtAB and its structure in the presence of ATP or AMP-PNP in this study, we also solved the structure of IrtAB_ΔSID_ in complex with ADP (Supplementary Figure 9 and Supplementary Table 1). As expected, this structure adopts a similar conformation to that of the ATP-free state (RMSD of 0.75 Å after superimposition of 1035 Cα atoms). Two obvious ADP densities are present in the NDBs of IrtA and IrtB (Supplementary Figure 9H). Therefore, this structure represents the state after ATP hydrolysis, but where ADP has not yet been released. Although the structural information presented here is not yet sufficient to reveal the mechanism of IrtAB mediated iron-loaded siderophores acquisition, it provides valuable insights for further studies to advance our understanding of its molecular mechanism.

A strong continuous non-protein density was found at the interface of IrtA and IrtB in the cryo-EM map. Specifically, this is between TMH2 in IrtA and TMH5 in IrtB, and the relative position and orientation of this density changed upon ATP binding (Supplementary Figure 8A, B). This density appears to belong to a partially ordered lipid molecule, implying that the lateral opening can accommodate hydrophobic acyl chains. Among the two types of siderophores with the same iron-binding core in *Mtb*, MBT with a lipid tail is considered to be anchored to the cell membrane (De Voss et al., 2000). Therefore, this portal is a likely route of entry for the hydrophobic acyl chain of MBT from the membrane into the binding cavity.

Although we have determined the structure of IrtAB in complex with ATP, it is still speculative as to whether the SID of IrtA affects the conformations of the TM region of IrtAB during ATP binding and hydrolysis. However, previous studies have shown that the SID of IrtA is not essential for Fe^3+^-cMBT import (Arnold et al., 2020). In addition, our ATPase activity assay also showed that this domain does not affect the ATP hydrolysis activities of IrtAB. Therefore, these results suggest that the SID of IrtA has little effect on the conformational changes in the TM region of IrtAB induced by ATP binding/hydrolysis. It has been shown that the ATPase activities of IrtAB can be strongly stimulated by the transport of the substrates, Fe^3+^-MBT and Fe^3+^-cMBT (Arnold et al., 2020). Combined with our structural data, the implication is that IrtAB forms an outward-facing conformation to expose a high-affinity binding site in the TM region, which could require its transport substrates.

ABC transporters play an extremely important role in transporting diverse substrates across the cell membrane. At present, the molecular mechanism as to how ABC exporter-like importers mediates substrate transport across the membrane is still largely unknown. The data presented here greatly advance our understanding of the molecular mechanism of this type of ABC importer. Importantly, there is an urgent need for the discovery of new anti-tuberculosis therapeutic drugs to deal with multi and extensive drug-resistant tuberculosis (Yang et al., 2020; Zhang et al., 2019), and siderophore analogues have already shown great potential as antibacterial agents (Juarez-Hernandez et al., 2012). Further understanding of the details of the IrtAB transport cycle, such as how substrate is recruited and released will be the next discovery that can contribute to our understanding of the functional mechanism of IrtAB, encouraging new efforts towards the discovery of novel tuberculosis drugs that target IrtAB.

## Materials and Methods

### Cloning and expression

The DNA coding sequences of full-length IrtAB (*Rv1348-Rv1349*) and the SID truncated IrtAB (IrtAB_ΔSID_) were amplified from *M. tuberculosis* H37Rv genomic DNA by PCR and cloned into a pM261 vector, fused with a C-terminal 10× His tag attached to IrtB. All mutations were generated using the Fast Mutagenesis System (TransGen). All constructs were verified by sequencing and then transformed into *M. smegmatis* mc^2^155 cells for expression. The cells were cultivated at 37°C in Luria Broth (LB) liquid media with shaking (220 rpm), supplemented with 50 μg mL^-1^ kanamycin, 20 μg mL^-1^ carbenicillin and 0.1% (v/v) Tween 80. The temperature was reduced to 16°C when the OD_600_ reached 1.0 and 0.2% (w/v) acetamide was added into the cell cultures to induce the overexpression of protein. After four days, cells were harvested by centrifugation at 4,000 rpm for 15 min and frozen at -80°C. All constructs were overexpressed using the same protocol as the wild-type protein.

### Protein purification

Full-length IrtAB, IrtAB_ΔSID_ and mutations were purified by a similar method. Cell pellets were thawed and resuspended in buffer A which contains 20 mM MES, 150 mM NaCl, pH 6.5. The resuspended cells were then lysed by a French Press at 1100 bar. Cell debris was removed by centrifugation at 12,000 rpm for 15 min at 4°C. The supernatant was collected and ultracentrifuged at 150,000 g for 1.5 h. The membrane fractions were collected and resuspended in buffer A. After incubating with 0.5% (w/v) LMNG (Anatrace) for 1.5 h at 4°C, the suspension was centrifuged and the supernatant was loaded onto a Ni-NTA agarose beads (Qiagen) affinity column supplemented with 10 mM imidazole. The beads were then washed in buffer A supplemented with 50 mM imidazole and 0.005% LMNG, and then exchanged into 0.06% digitonin buffer. The protein was eluted from the beads with buffer A supplemented with 500 mM imidazole and 0.06% digitonin, then concentrated and loaded to a size exclusion chromatography column (Superose 6 Increase 10/300 GL, GE Healthcare) pre-equilibrated with 20 mM MES, 150 mM NaCl, 2 mM DTT and 0.06% digitonin, pH 6.5. The peak fractions were pooled and concentrated for cryo-EM sample preparation using a 100-kDa cut-off concentrator (Merck Millipore). The results of protein purification are shown in Fig. S1.

### ATPase activity assay

The ATPase Activity Assay Kit (Sigma-Aldrich, MAK-113) was used to measure ATPase activity. The protein was diluted by the buffer (MAK-113) to a final concentration of 2 μM, incubated with 1 mM ATP in 40 μL reaction mixture which contained 20 mM MES pH 6.5, 150 mM NaCl, 0.06% digitonin, 5 mM MgCl_2_ for 30 min at 37°C. Next, the enzyme reactions were stopped and the colorimetric product was generated by adding 200 μL of Reagent (MAK-113) into each reaction well. After incubating for an additional 30 minutes at room temperature, the absorbance at 620 nm in each well was measured. ATPase activity was represented as the amount of phosphate produced from the ATP catalytic reaction. The experiments were performed in triplicate. Results were analyzed in GraphPad Prism 5.0 (https://www.graphpad.com).

### Cryo-EM sample preparation

The protein samples were concentrated to 5 mg ml^-1^. For IrtAB_ΔSID_, 1 mM adenosine 5′-(β,γ-imido)triphosphate (AMP-PNP) and MgCl_2_ or 5 mM ADP-MgCl_2_ and 5 mM DTT were added and incubated on ice for 30 min. For mutated IrtAB (E-Q) and IrtAB_ΔSID_ (E-Q), 10mM ATP-MgCl_2_ and 5 mM DTT were added and incubated on ice for 30 min. All protein samples were combined with various ligands and then centrifuged before preparation for freezing samples. 3 μl of protein sample was applied to holey carbon grids (Quantifoil, R 0.6/1 Cu 200 mesh) which were H_2_/O_2_ glow-discharged for 30 s. After blotting for 3 s, the grids were frozen in liquid ethane using an FEI Vitrobot Mark IV at 8°C and 100% humidity, and then transferred into liquid nitrogen.

### Cryo-EM data collection

For the full-length IrtAB sample, cryo-EM data were collected on a 300 kV FEI Titan Krios electron microscope with a Gatan K3 Summit direct electron detector at a nominal magnification of 29,000 with a pixel size of 0.82 Å in super-resolution mode. Movies were recorded for 2.4 s in 40 sub-frames with a total dose of 60 e^-^/Å^2^. For other protein samples, a Gatan K2 Summit direct electron detector, equipped with a Gatan quantum energy filter, were used to capture movies at EFTEM magnification of 165,000 in super-resolution mode with a pixel size of 0.82 Å.

### Cryo-EM image processing and 3D Reconstruction

For the image processing of full-length IrtAB, a total of 1875 original movies were generated and beam-induced correction was performed by MotionCor2 (Zheng et al., 2017), the remaining steps for image processing were performed using cryoSPARC (Punjani et al., 2017). After CTF estimation, poor-quality images with features such as high astigmatism, low CTF fit resolution and large-offset defocus values were manually removed. 428,375 particles were automatically selected and extracted with a box size of 384 pixels from selected images. Several rounds of 2D classification were performed to remove bad particles, and then *Ab-initio* reconstruction was performed with 308,525 particles to generate 3D volumes as templates for heterogeneous refinement, with 143,181 particles converged into one good class. Homogeneous refinement was performed on this convergence class, followed by non-uniform refinement and local refinement. Finally, a density map was obtained with an estimated average resolution of 3.48 Å according to the gold-standard Fourier shell correlation (FSC) cut-off of 0.143 (Grigorieff, 2016). Local resolution ranges were analyzed within cryoSPARC. The other datasets were processed in the same way.

### Model building and refinement

The model of full-length IrtAB was manually built, but relied on the 3.5-Å cryo-EM map and the crystal structure of truncated *Mycobacterium thermoresistibile* IrtAB (PDB:6TEG) (Arnold et al., 2020). This model was docked into the cryo-EM map in UCSF Chimera (Pettersen et al., 2004) and manually adjusted in Coot (Emsley et al., 2010). Several iterations of real-space refinement was performed in PHENIX (Adams et al., 2010). The models for other states of IrtAB complex were generated based on their cryo-EM maps and using the model of full-length IrtAB as a starting point. Nucleotides were fitted as rigid-bodies into the cryo-EM map using Coot. The structures were refined in real space using PHENIX with secondary structure and geometry restraints in place to prevent overfitting. The final atomic model was evaluated using MolProbity (Chen et al., 2010). Cryo-EM data collection and model refinement statistics are shown in Appendix Table S1.

All graphics were generated using PyMOL (http://www.pymol.org), UCSF ChimeraX (Goddard et al., 2018), UCSF Chimera.

## Acknowledgements

We thank the staff from the Electron Microscopy Facility of ShanghaiTech University, for assistance during cryo-EM data collection. We are thankful to the Analytical chemistry platform of Shanghai Institute for Advanced Immunochemical studies (SIAIS) for their assistance in mass spectrometry analysis.

## Funding

This work was supported by grants from the National Key Research and Development Program of China (Grant No. 2017YFC0840300) and the National Natural Science Foundation of China (Grant No. 81520108019 to R.Z.; 81772204 to H.Y.; 32171217 to B.Z.) and the Linggang Laboratory (Grant No. LG202101-01-08).

## Author contributions

Z.R., B.Z. and H.Y. initiated and supervised the project. B.Z. and S.S. designed experiments. S.S. made all constructs and purified the proteins. S.S. and Y.G. collected and processed cryo-EM data. B.Z. and S.S. built and refined the structure models. S.S., Xiaolin Yang performed functional experiments with help from Xiuna Yang, T.H., J.L., Z.X., P.R. and F.B.. S.S., B.Z., Xiaolin Yang, Xiuna Yang, T.H., J.L., Z.X., H.Y. and Z.R. analyzed and discussed the results. The manuscript was written by B.Z., S.S., H.Y., L.W.G. and Z.R. with the help of all the authors.

## Competing interest declaration

All authors declare that they have no conflict of interest.

## Data and materials availability

All data are available in the manuscript. The accession number for the 3D cryo-EM density maps reported in this paper is EMD-32536, EMD-32537, EMD-32538 and EMD-32539. Atomic coordinates and structure factors for the IrtAB, IrtAB_ΔSID_-AMP-PNP, IrtAB_ΔSID_ (E-Q)-ATP and IrtAB_ΔSID_-ADP structures have been deposited in the Protein Data Bank with identification codes 7WIU, 7WIV, 7WIW and 7WIX.

## Figures and Legends

**Supplementary Figure 1.**
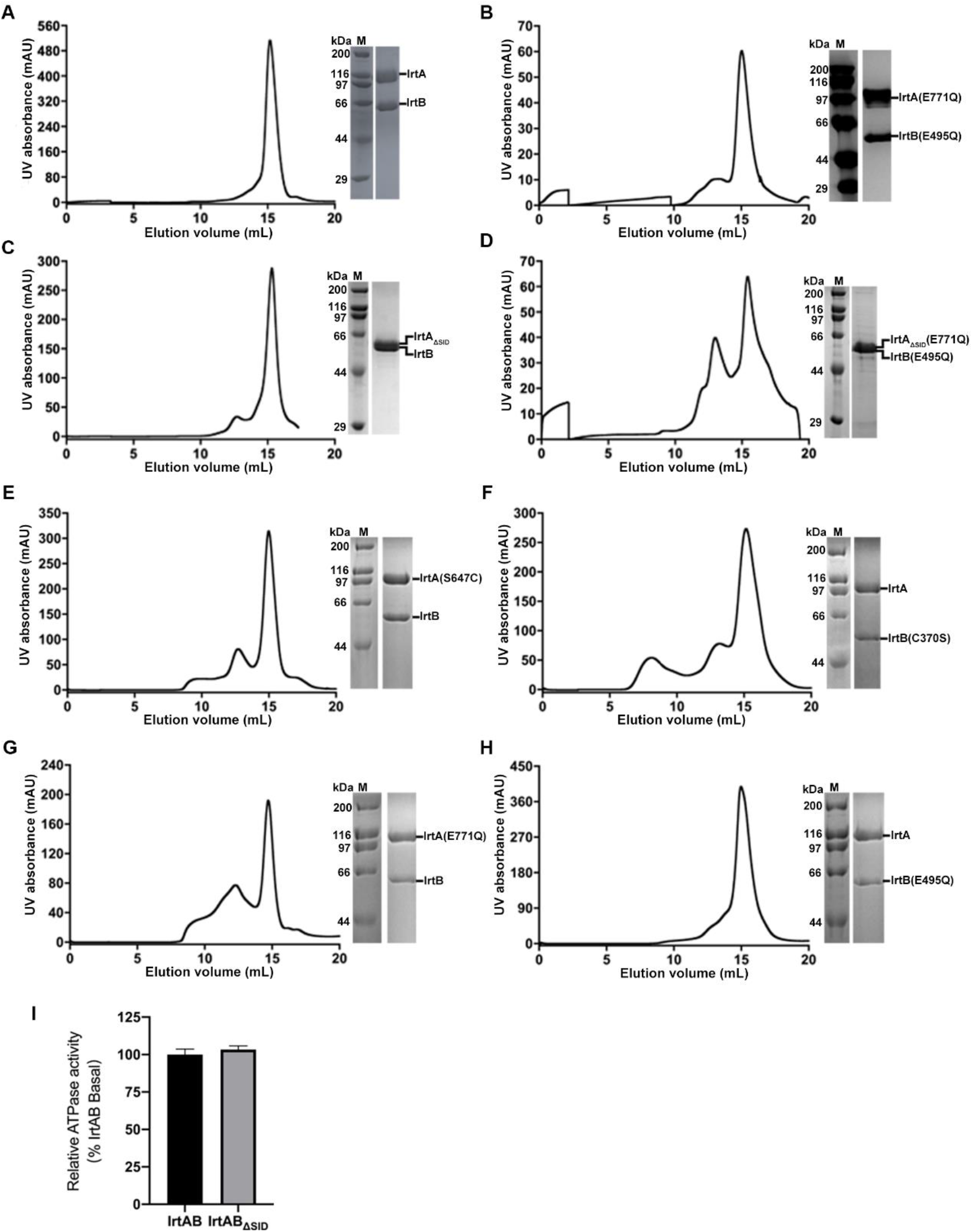
Properties of *M. tuberculosis* IrtAB. (**A**-**H**) Size exclusion profile and SDS-PAGE of the purified different samples in digitonin. (**I**) ATPase activity of SID truncated IrtAB (IrtAB_ΔSID_) was normalized relative to that of the full-length IrtAB. Error bars represent mean ± SD based on three independent measurements.

**Supplementary Figure 2.**
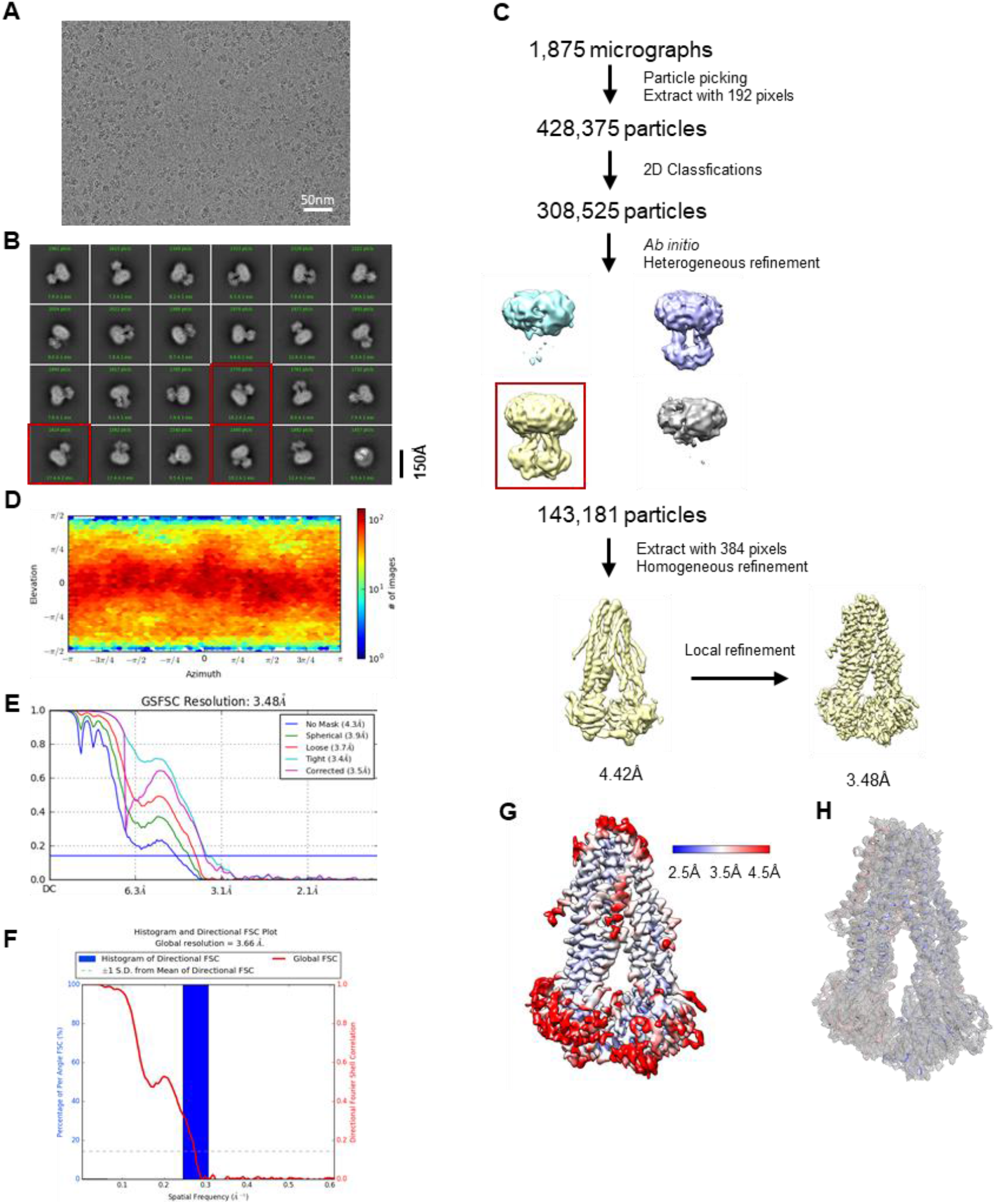
Cryo-EM data processing of the ATP-free, full-length IrtAB. (**A**) Representative cryo-EM image of the full-length IrtAB complex. (**B**) Representative 2D classification averages showing the full-length IrtAB in different orientations. The SID of IrtA is observed in some views indicated by red box. (**C**) Summary of the image processing procedure. (**D**) Angular distribution heatmap of particles used for the refinement. Each sphere indicates the number of particles from this angle, and the size of the spheres corresponds to the number of particles. (**E**) Fourier shell correlation (FSC) curves of the final 3D reconstruction. (gold-standard FSC curve between the two half maps with indicated resolution at FSC=0.143). (**F**) 3D FSC histogram of the final map. (**G**) Local resolution of the final cryo-EM map of the full-length IrtAB. (**H**) Cryo-EM map density (gray mesh, contoured at 7 σ) for the structure of full-length IrtAB without nucleotide bound.

**Supplementary Figure 3.**
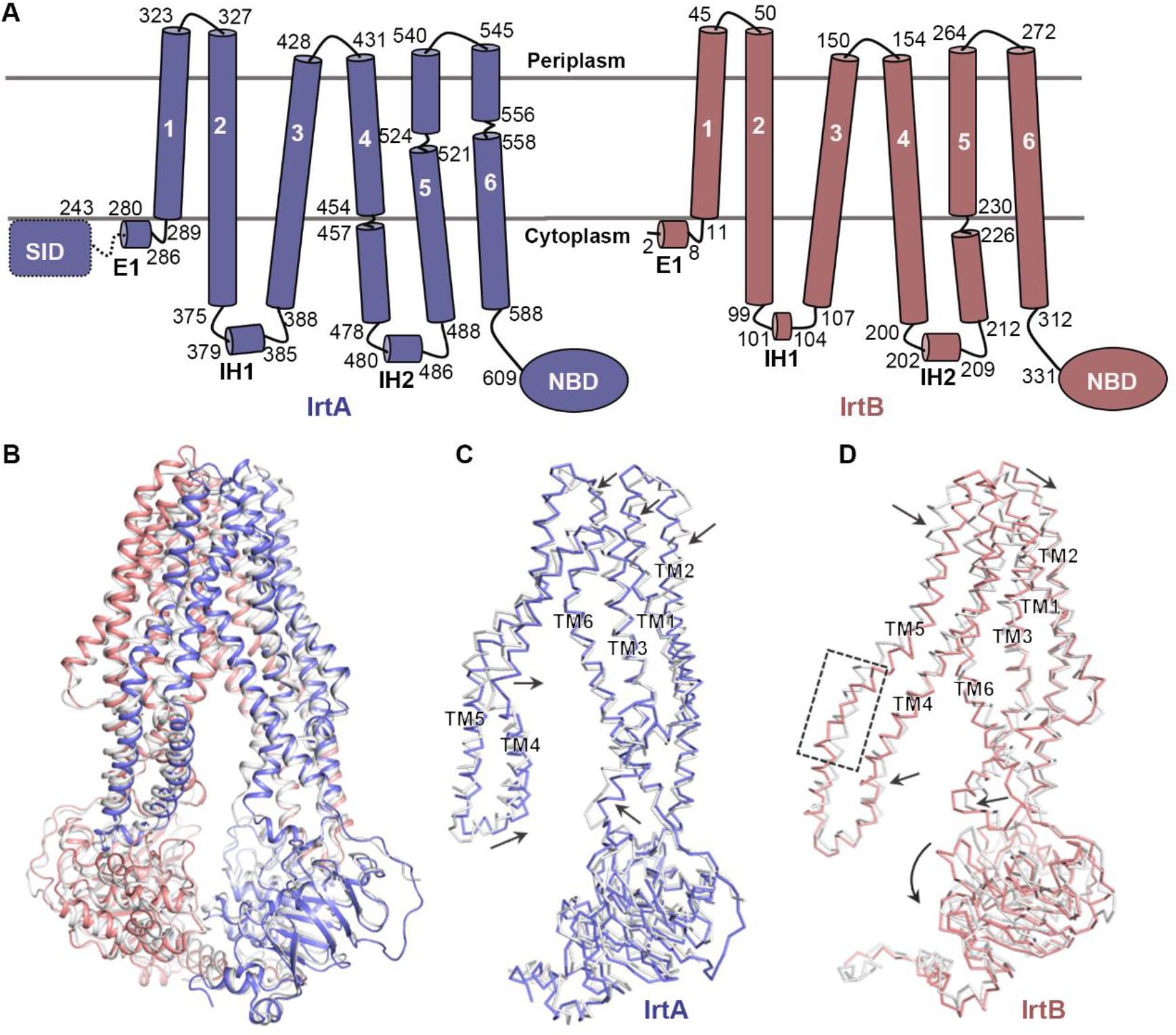
Structural comparison of IrtAB in mycobacteria. (**A**) Topology diagram of full-length IrtA (slate) and IrtB (salmon) with TMHs (transmembrane helices) numbered. The amino acid sequence numbers are indicated. The linkage between the siderophore interaction domain (SID) (dotted frame) and elbow helices (E1) is indicated by the dotted lines. IH, intracellular helix. NBD, nucleotide-binding domain. (**B**) Superposition of *Mtb* IrtAB (IrtA, slate; IrtB, salmon) and *Mycobacterium thermoresistibile* IrtAB (PDB: 6TEJ, white) structures. (**C**) Superimposition of IrtA structures in *Mtb* (slate) and *Mycobacterium thermoresistibile* (white). The divergence in these two structures is indicated by the black arrows. (**D**) Superimposition of IrtB structures in *Mtb* (salmon) and *Mycobacterium thermoresistibile* (white). The differences between these two structures are indicated by black arrows and a dashed box.

**Supplementary Figure 4.**
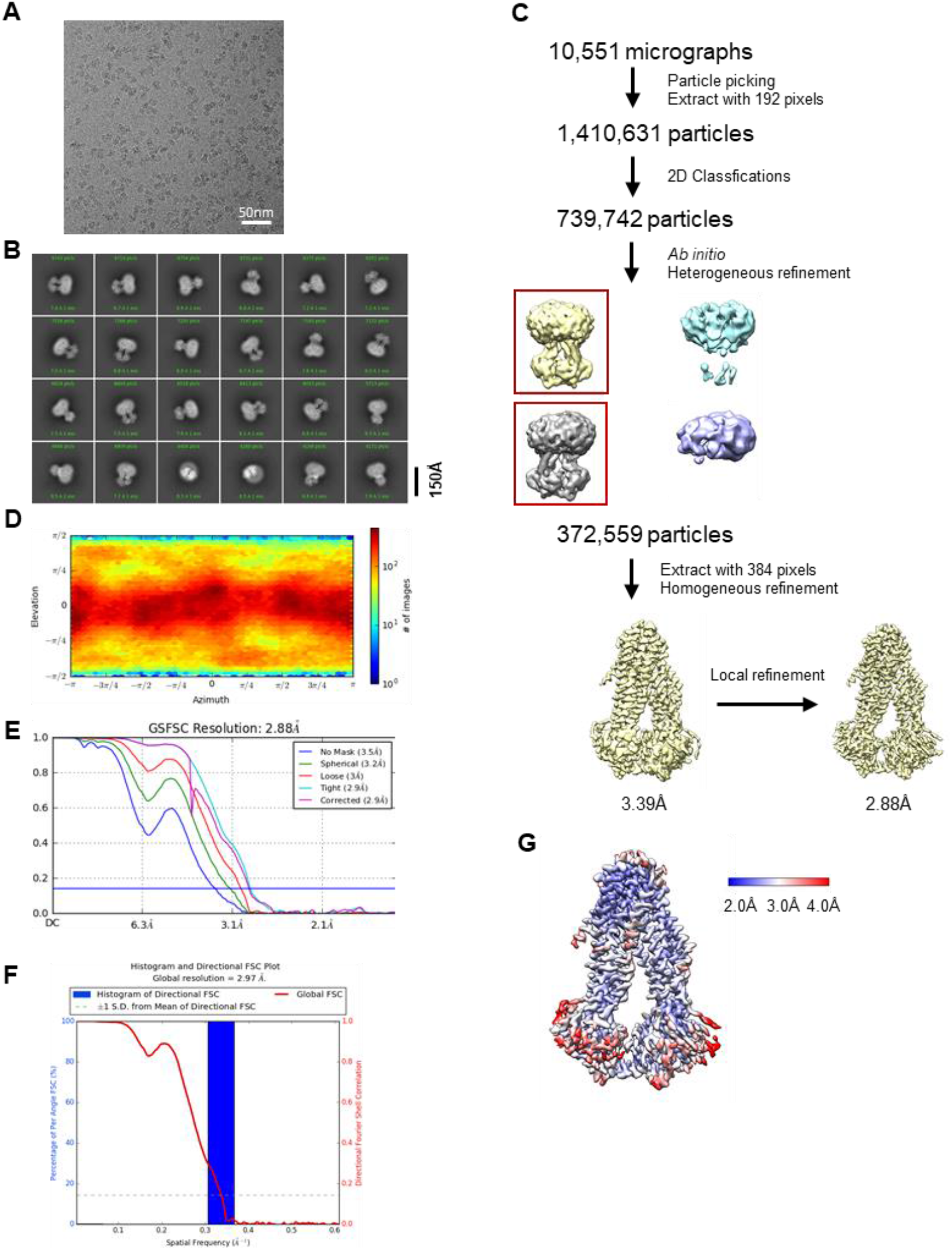
Cryo-EM data processing of the AMP-PNP-bound, SID truncated IrtAB. (**A**) Representative cryo-EM image of IrtAB_ΔSID_ in complex with AMP-PNP. (**B**) Representative 2D classification averages showing the complex in different orientations. (**C**) Summary of the image processing procedure. (**D**) Angular distribution heatmap of particles used for the refinement. (**E**) Fourier shell correlation (FSC) curves of the final 3D reconstruction. (**F**) 3D FSC histogram of the final map. (**G**) Local resolution of the final cryo-EM map of IrtAB_ΔSID_ in complex with AMP-PNP.

**Supplementary Figure 5.**
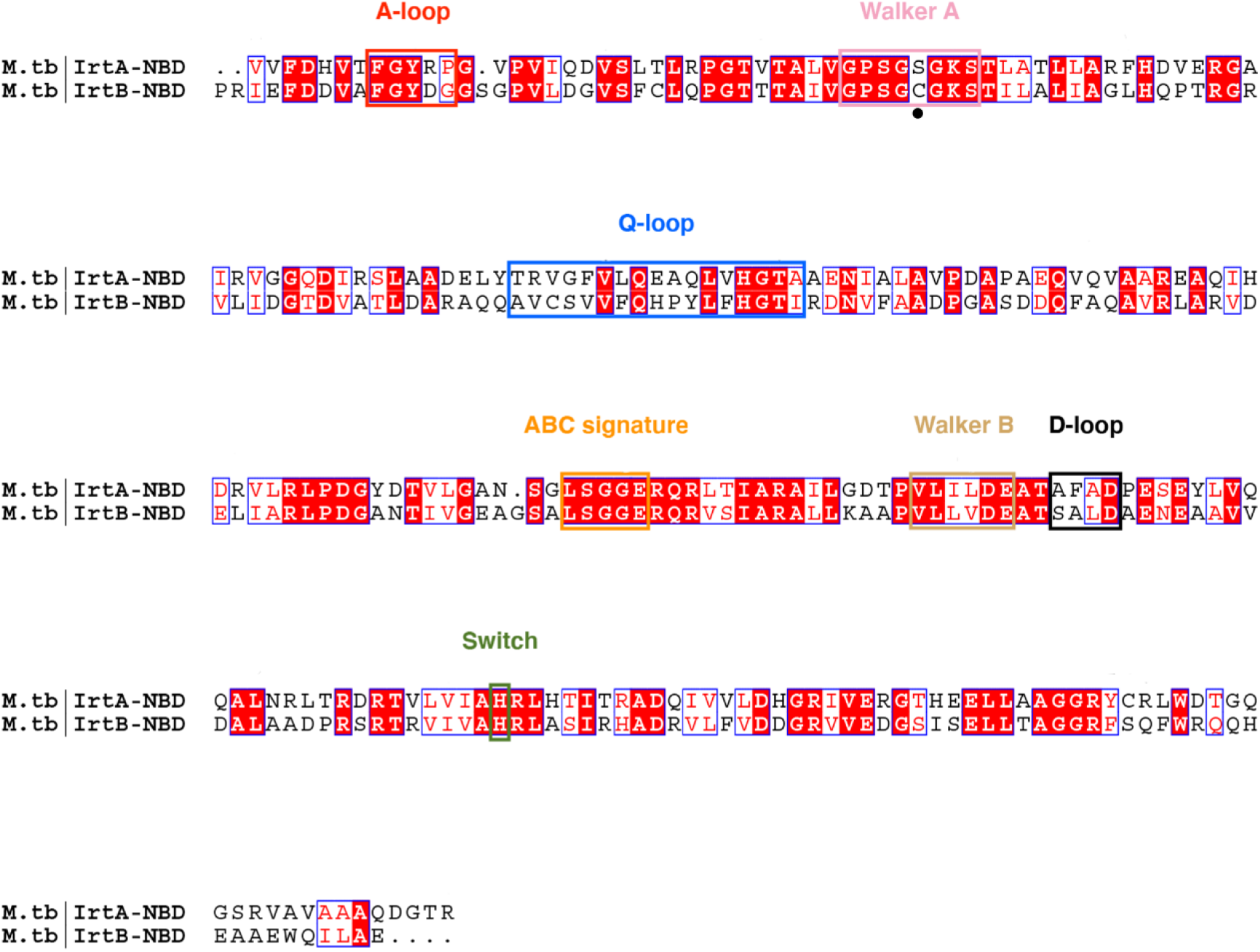
Sequence alignment of NBDs of IrtA and IrtB. The conserved sequence motifs of NBD are also indicated.

**Supplementary Figure 6.**
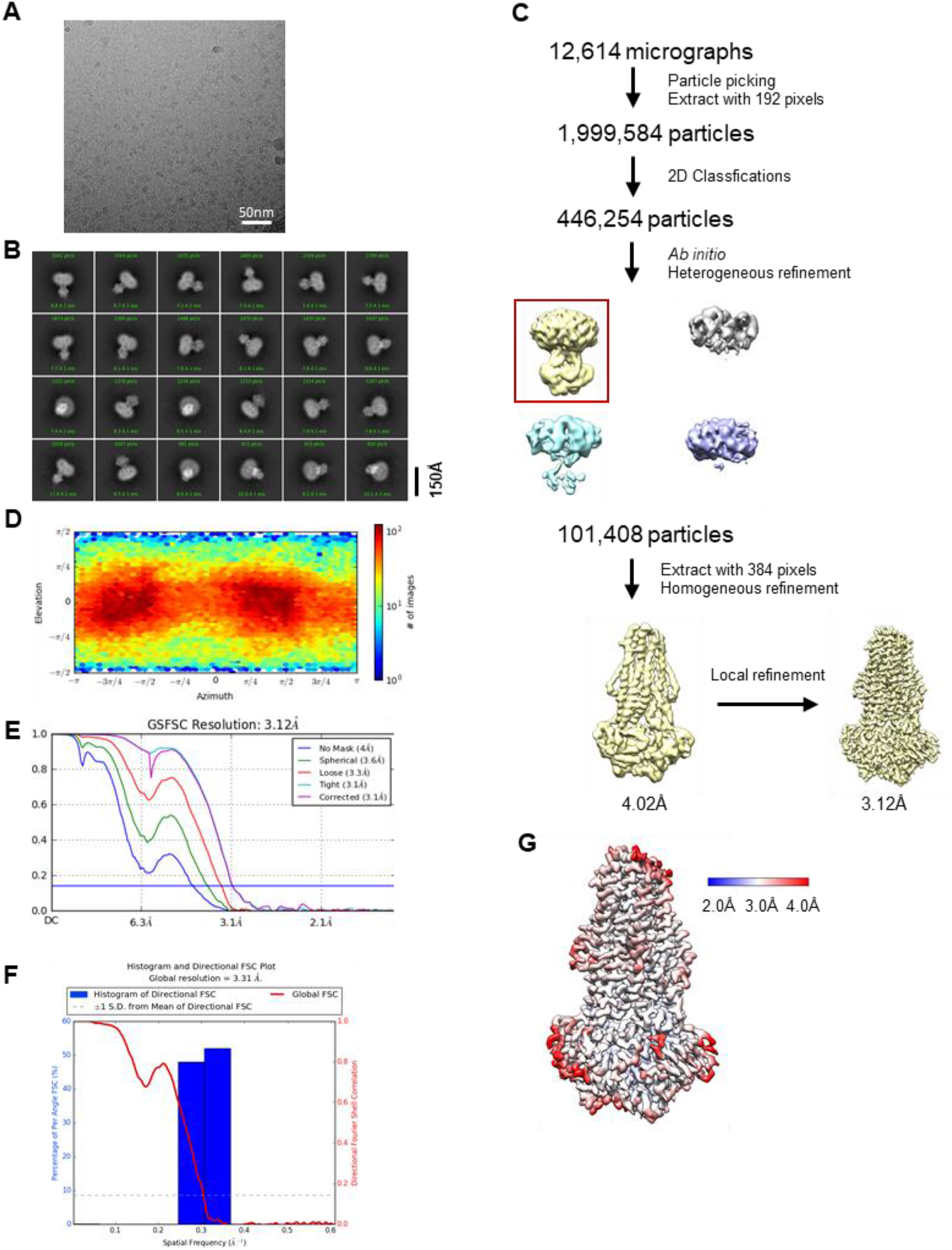
Cryo-EM data processing of IrtAB_ΔSID_ (E-Q) bound ATP. (**A**) Representative cryo-EM image of IrtAB_ΔSID_ (E-Q) in complex with ATP. (**B**) Representative 2D classification averages showing the complex in different orientations. (**C**) Summary of the image processing procedure. (**D**) Angular distribution heatmap of particles used for the refinement. (**E**) Fourier shell correlation (FSC) curves of the final 3D reconstruction. (**F**) 3D FSC histogram of the final map. (**G**) Local resolution of the final cryo-EM map of IrtAB_ΔSID_ (E-Q) in complex with ATP.

**Supplementary Figure 7.**
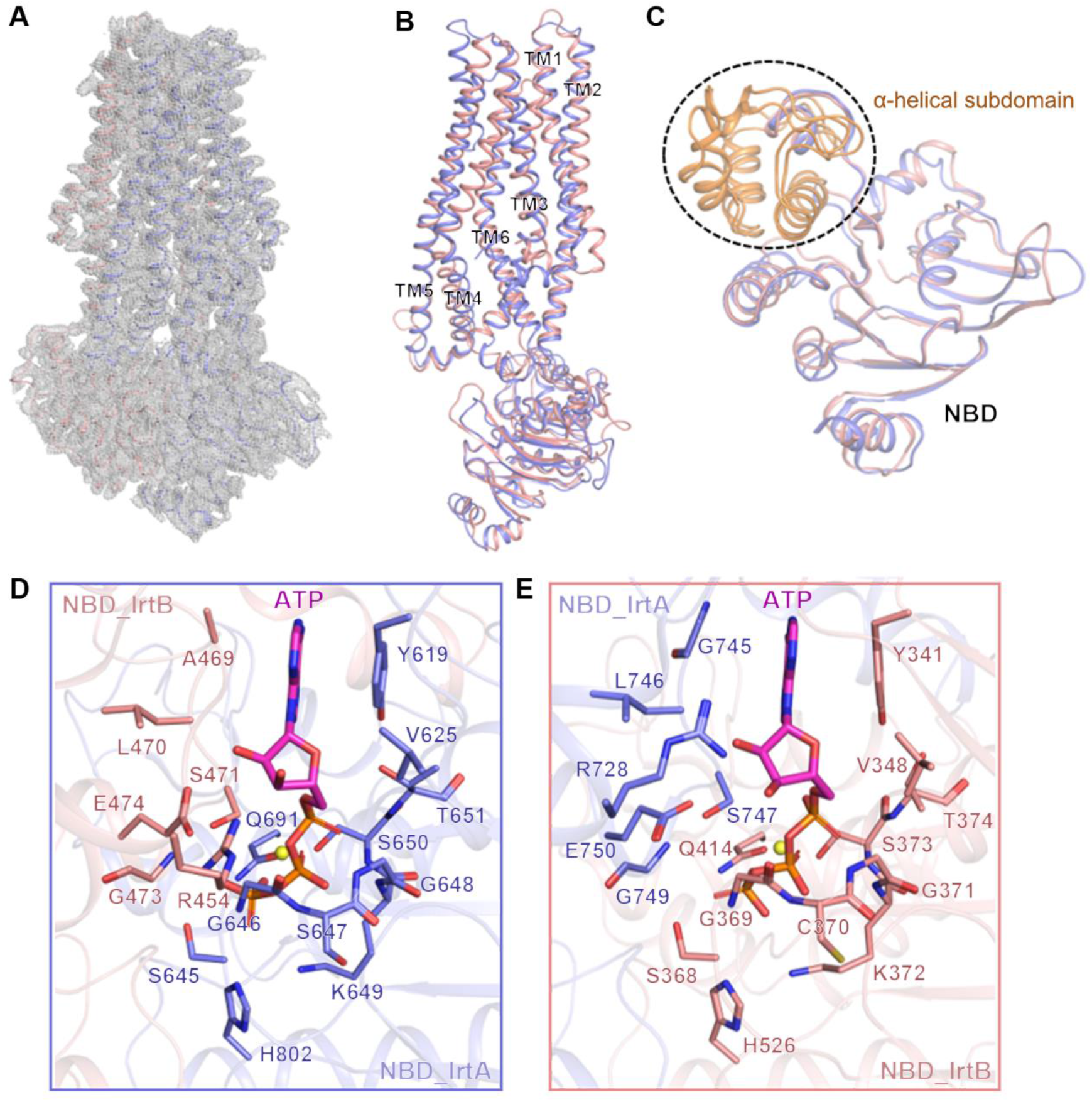
Structural analysis of IrtAB_ΔSID_ (E-Q) in complex with ATP. (**A**) Cryo-EM map density (gray mesh, contoured at 9 σ) for the structure of IrtAB_ΔSID_ (E-Q) in complex with ATP. (**B**) Superimposition of IrtA (slate) and IrtB (salmon) structures. (**C**) Superimposition of NBD in IrtA (slate) and NBD in IrtB (salmon). The α-helical subdomain is colored orange and indicated with a dotted line. (**C**) Close-up view of the ATPase site in IrtA. The residues involved in ATP binding are shown as sticks. ATP is represented by magenta sticks colored by heteroatom. The Mg^2+^ is shown as a yellow sphere. (**E**) Close-up view of the ATPase site in IrtB. The residues involved in ATP binding are shown as sticks. ATP is represented by magenta sticks colored by heteroatom. The Mg^2+^ is shown as a yellow sphere.

**Supplementary Figure 8.**
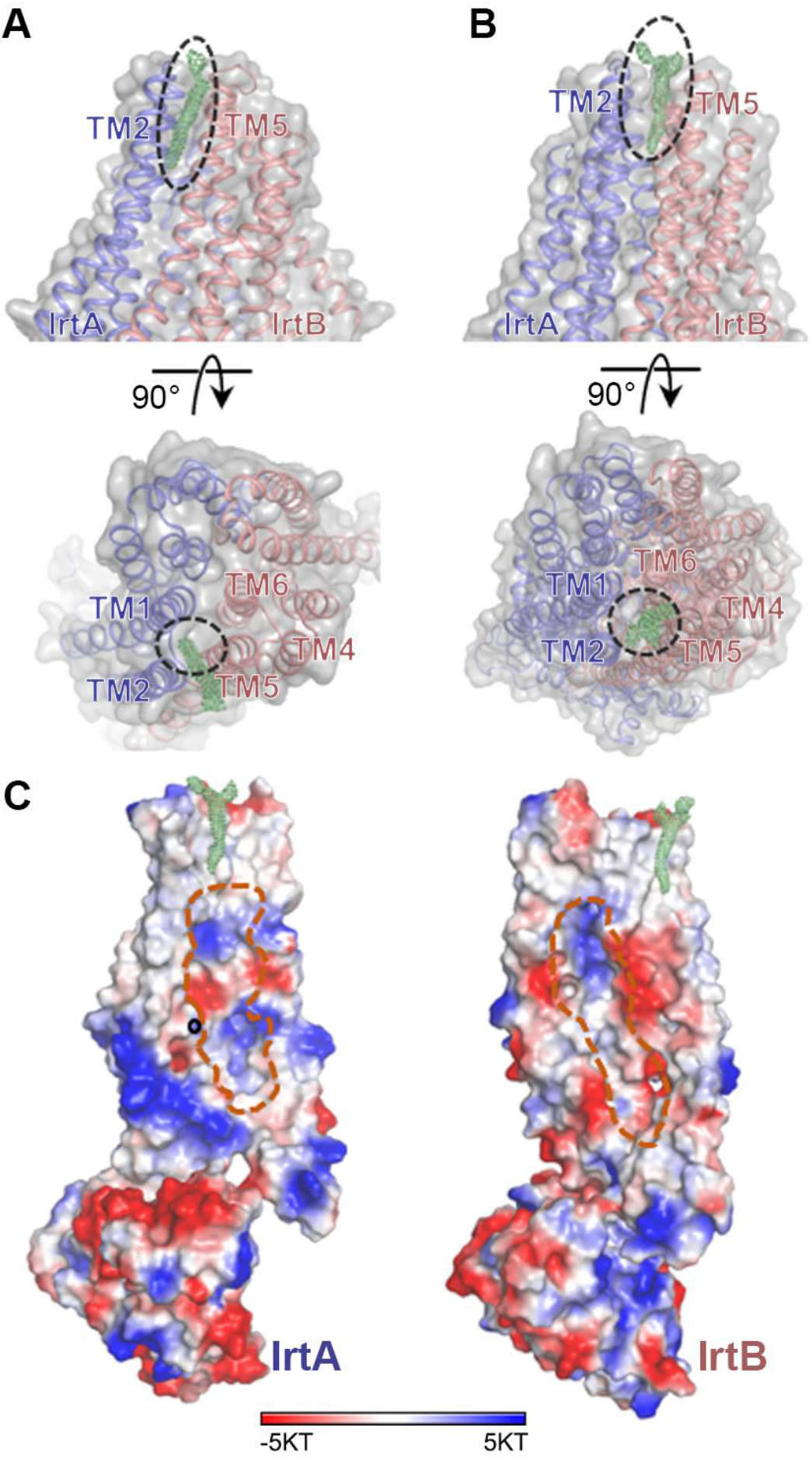
The portal between the interface of IrtA and IrtB. (**A**) Two orthogonal views of the TM region interface between IrtA and IrtB in the open-inward conformation. The portal between TMH2 of IrtA and TMH5 of IrtB is indicated with a dotted line. Non-protein cryo-EM map density with the lateral opening contoured at 7 σ is shown as green mesh. (**B**) Two orthogonal views of the TM region interface between IrtA and IrtB in the occluded conformation. Non-protein cryo-EM map density surrounded by TMH1-2 of IrtA and TMH5-6 of IrtB contoured at 7 σ is shown as a green mesh. (**C**) The electrostatic inner surface representation of IrtA and IrtB in the occluded conformation. The cavity is outlined with a broken orange line. The surfaces are colored from blue (most positive) to red (most negative). Hydrophobic regions are shown in white. Non-protein cryo-EM map density is also indicated with a green mesh. The binding site of the metal ion in IrtA is indicated with a black circle.

**Supplementary Figure 9.**
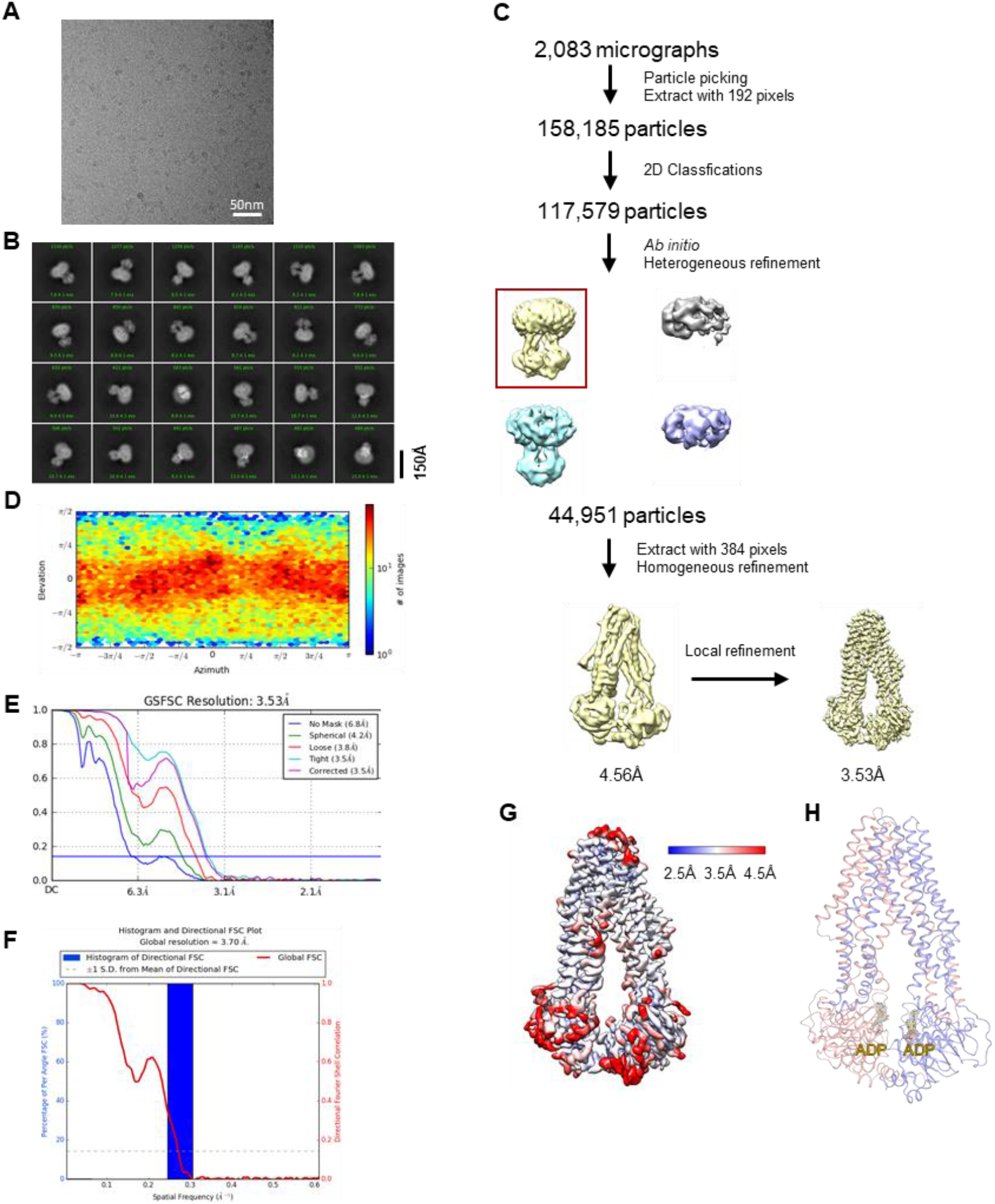
Cryo-EM data processing of the ADP-bound, SID truncated IrtAB (IrtAB_ΔSID_). (**A**) Representative cryo-EM image of IrtAB_ΔSID_ in complex with ADP. (**B**) Representative 2D classification averages showing the complex in different orientations. (**C**) Summary of the image processing procedure. (**D**) Angular distribution heatmap of particles used for the refinement. (**E**) Fourier shell correlation (FSC) curves of the final 3D reconstruction. (**F**) 3D FSC histogram of the final map. (**G**) Local resolution of the final cryo-EM map of IrtAB_ΔSID_ in complex with ADP. (**H**) Cartoon representation of the structure of IrtAB_ΔSID_ in complex with ADP. The density map for ADP (yellow sticks) is shown as gray mesh and contoured at 9 σ.

**Supplementary Table 1.** Statistics for the cyro-EM structures presented in this study.

|  | IrtAB | IrtAB<sub>ΔSID</sub>-AMP-PNP | IrtAB<sub>ΔSID</sub> (E-Q)-ATP | IrtAB<sub>ΔSID</sub>-ADP |
| --- | --- | --- | --- | --- |
| Microscope | FEI Titan Krios |  |  |  |
| Magnification | 29,000 x | 165,000 x |  |  |
| Voltage (keV) | 300 |  |  |  |
| Electron exposure (e<sup>-</sup>/Å<sup>2</sup>) | 60 |  |  |  |
| Defocus range (μm) | -1.2 to -1.8 |  |  |  |
| Pixel size (Å/pixel) | 0.82 |  |  |  |
| Number of movies | 1,875 | 10,551 | 12,614 | 2,083 |
| Symmetry imposed | C1 |  |  |  |
| Final particle images (no.) | 143,181 | 372,559 | 101,408 | 44,951 |
| Map resolution (Å) | 3.48 | 2.88 | 3.12 | 3.53 |
| FSC threshold | 0.143 | 0.143 | 0.143 | 0.143 |
| Map resolution range (Å) | 3.0 – 11.8 | 2.4 – 9.8 | 2.7 – 11.8 | 3.0 – 13.7 |
| **Refinement** |  |  |  |  |
| Initial model used (PDB code) | 6TEJ |  |  |  |
| Model resolution (Å) | 3.48 | 2.88 | 3.12 | 3.53 |
| FSC threshold | 0.143 | 0.143 | 0.143 | 0.143 |
| Model resolution range (Å) | 3.48 | 2.88 | 3.12 | 3.53 |
| Map sharpening *B* factor (Å<sup>2</sup>) | -69.4 | -69.4 | -77.0 | -64.3 |
| Model composition |  |  |  |  |
| Non-hydrogen atoms | 8615 | 8646 | 8603 | 8670 |
| Protein residues | 1147 | 1147 | 1137 | 1147 |
| Ligands | - | 1 | 5 | 3 |
| *B* factors (Å<sup>2</sup>) |  |  |  |  |
| Protein | 99.24 | 57.88 | 52.33 | 95.56 |
| Ligand | - | 70.48 | 45.37 | 100.23 |
| R.m.s. deviations |  |  |  |  |
| Bond lengths (Å) | 0.003 | 0.013 | 0.004 | 0.010 |
| Bond angles (°) | 0.814 | 0.973 | 0.878 | 1.029 |
| Validation |  |  |  |  |
| MolProbity score | 2.00 | 1.96 | 1.87 | 2.22 |
| Clashscore | 7.61 | 7.19 | 7.05 | 11.27 |
| Poor rotamers (%) | 0.00 | 0.68 | 0.00 | 0.23 |
| Ramachandran plot |  |  |  |  |
| Favored (%) | 88.89 | 89.41 | 92.06 | 87.66 |
| Allowed (%) | 11.11 | 10.15 | 7.59 | 11.90 |
| Disallowed (%) | 0.00 | 0.44 | 0.35 | 0.44 |

## Notes

### Competing Interest Statement

The authors have declared no competing interest.

## References

Adams, P.D., Afonine, P.V., Bunkoczi, G., Chen, V.B., Davis, I.W., Echols, N., Headd, J.J., Hung, L.W., Kapral, G.J., Grosse-Kunstleve, R.W., McCoy, A.J., Moriarty, N.W., Oeffner, R., Read, R.J., Richardson, D.C., Richardson, J.S., Terwilliger, T.C., Zwart, P.H., 2010. PHENIX: a comprehensive Python-based system for macromolecular structure solution. Acta Crystallogr D Biol Crystallogr 66: 213–221. DOI: https://doi.org/10.1107/S0907444909052925, PMID: 20124702

Arnold, F.M., Weber, M.S., Gonda, I., Gallenito, M.J., Adenau, S., Egloff, P., Zimmermann, I., Hutter, C.A.J., Hurlimann, L.M., Peters, E.E., Piel, J., Meloni, G., Medalia, O., Seeger, M.A., 2020. The ABC exporter IrtAB imports and reduces mycobacterial siderophores. Nature 580: 413–417. DOI: https://doi.org/10.1038/s41586-0202136-9, PMID: 32296173

Arosio, P., Elia, L., Poli, M., 2017. Ferritin, cellular iron storage and regulation. IUBMB Life 69: 414–422. DOI: https://doi.org/10.1002/iub.1621, PMID: 28349628

Begg, S.L., 2019. The role of metal ions in the virulence and viability of bacterial pathogens. Biochem Soc Trans 47: 77–87. DOI: https://doi.org/10.1042/BST20180275, PMID: 30626704

Braibant, M., Gilot, P., Content, J., 2000. The ATP binding cassette (ABC) transport systems of Mycobacterium tuberculosis. FEMS Microbiol Rev 24: 449–467. DOI: https://doi.org/10.1111/j.1574-6976.2000.tb00550.x, PMID: 10978546

Brock, J.H., 2012. Lactoferrin--50 years on. Biochem Cell Biol 90: 245–251. DOI: https://doi.org/10.1139/o2012-018, PMID: 22574842

Chen, J., 2013. Molecular mechanism of the Escherichia coli maltose transporter. Curr Opin Struct Biol 23: 492–498. DOI: https://doi.org/10.1016/j.sbi.2013.03.011, PMID: 23628288

Chen, V.B., Arendall, W.B., 3rd, Headd, J.J., Keedy, D.A., Immormino, R.M., Kapral, G.J., Murray, L.W., Richardson, J.S., Richardson, D.C., 2010. MolProbity: all-atom structure validation for macromolecular crystallography. Acta Crystallogr D Biol Crystallogr 66: 12–21. DOI: https://doi.org/10.1107/S0907444909042073, PMID: 20057044

Choudhury, H.G., Tong, Z., Mathavan, I., Li, Y., Iwata, S., Zirah, S., Rebuffat, S., van Veen, H.W., Beis, K., 2014. Structure of an antibacterial peptide ATP-binding cassette transporter in a novel outward occluded state. Proc Natl Acad Sci U S A 111: 9145–9150. DOI: https://doi.org/10.1073/pnas.1320506111, PMID: 24920594

Dawson, R.J., Locher, K.P., 2006. Structure of a bacterial multidrug ABC transporter. Nature 443: 180–185. DOI: https://doi.org/10.1038/nature05155, PMID: 16943773

de Jong, G., van Dijk, J.P., van Eijk, H.G., 1990. The biology of transferrin. Clin Chim Acta 190: 1–46. DOI: https://doi.org/10.1016/0009-8981(90)90278-z, PMID: 2208733

De Voss, J.J., Rutter, K., Schroeder, B.G., Su, H., Zhu, Y., Barry, C.E., 3rd, 2000. The salicylate-derived mycobactin siderophores of Mycobacterium tuberculosis are essential for growth in macrophages. Proc Natl Acad Sci U S A 97: 1252–1257. DOI: https://doi.org/10.1073/pnas.97.3.1252, PMID: 10655517

Emsley, P., Lohkamp, B., Scott, W.G., Cowtan, K., 2010. Features and development of Coot. Acta Crystallogr D Biol Crystallogr 66: 486–501. DOI: https://doi.org/10.1107/S0907444910007493, PMID: 20383002

Esser, L., Zhou, F., Pluchino, K.M., Shiloach, J., Ma, J., Tang, W.K., Gutierrez, C., Zhang, A., Shukla, S., Madigan, J.P., Zhou, T., Kwong, P.D., Ambudkar, S.V., Gottesman, M.M., Xia, D., 2017. Structures of the Multidrug Transporter P-glycoprotein Reveal Asymmetric ATP Binding and the Mechanism of Polyspecificity. J Biol Chem 292: 446–461. DOI: https://doi.org/10.1074/jbc.M116.755884, PMID: 27864369

Gobin, J., Moore, C.H., Reeve, J.R., Jr., Wong, D.K., Gibson, B.W., Horwitz, M.A., 1995. Iron acquisition by Mycobacterium tuberculosis: isolation and characterization of a family of iron-binding exochelins. Proc Natl Acad Sci U S A 92: 5189–5193. DOI: https://doi.org/10.1073/pnas.92.11.5189, PMID: 7761471

Goddard, T.D., Huang, C.C., Meng, E.C., Pettersen, E.F., Couch, G.S., Morris, J.H., Ferrin, T.E., 2018. UCSF ChimeraX: Meeting modern challenges in visualization and analysis. Protein Sci 27: 14–25. DOI: https://doi.org/10.1002/pro.3235, PMID: 28710774

Grigorieff, N., 2016. Frealign: An Exploratory Tool for Single-Particle Cryo-EM. Methods Enzymol 579: 191–226. DOI: https://doi.org/10.1016/bs.mie.2016.04.013, PMID: 27572728

Hofmann, S., Januliene, D., Mehdipour, A.R., Thomas, C., Stefan, E., Bruchert, S., Kuhn, B.T., Geertsma, E.R., Hummer, G., Tampe, R., Moeller, A., 2019. Conformation space of a heterodimeric ABC exporter under turnover conditions. Nature 571: 580–583. DOI: https://doi.org/10.1038/s41586-019-1391-0, PMID: 31316210

Hohl, M., Briand, C., Grutter, M.G., Seeger, M.A., 2012. Crystal structure of a heterodimeric ABC transporter in its inward-facing conformation. Nat Struct Mol Biol 19: 395–402. DOI: https://doi.org/10.1038/nsmb.2267, PMID: 22447242

Jin, M.S., Oldham, M.L., Zhang, Q., Chen, J., 2012. Crystal structure of the multidrug transporter P-glycoprotein from Caenorhabditis elegans. Nature 490: 566–569. DOI: https://doi.org/10.1038/nature11448, PMID: 23000902

Juarez-Hernandez, R.E., Franzblau, S.G., Miller, M.J., 2012. Syntheses of mycobactin analogs as potent and selective inhibitors of Mycobacterium tuberculosis. Org Biomol Chem 10: 7584–7593. DOI: https://doi.org/10.1039/c2ob26077h, PMID: 22895786

Khare, D., Oldham, M.L., Orelle, C., Davidson, A.L., Chen, J., 2009. Alternating access in maltose transporter mediated by rigid-body rotations. Mol Cell 33: 528–536. DOI: https://doi.org/10.1016/j.molcel.2009.01.035, PMID: 19250913

Kim, Y., Chen, J., 2018. Molecular structure of human P-glycoprotein in the ATP-bound, outward-facing conformation. Science 359: 915–919. DOI: https://doi.org/10.1126/science.aar7389, PMID: 29371429

Kodan, A., Yamaguchi, T., Nakatsu, T., Sakiyama, K., Hipolito, C.J., Fujioka, A., Hirokane, R., Ikeguchi, K., Watanabe, B., Hiratake, J., Kimura, Y., Suga, H., Ueda, K., Kato, H., 2014. Structural basis for gating mechanisms of a eukaryotic P-glycoprotein homolog. Proc Natl Acad Sci U S A 111: 4049–4054. DOI: https://doi.org/10.1073/pnas.1321562111, PMID: 24591620

Liu, F., Liang, J., Zhang, B., Gao, Y., Yang, X., Hu, T., Yang, H., Xu, W., Guddat, L.W., Rao, Z., 2020. Structural basis of trehalose recycling by the ABC transporter LpqY-SugABC. Sci Adv 6. DOI: https://doi.org/10.1126/sciadv.abb9833, PMID: 33127676

Locher, K.P., 2016. Mechanistic diversity in ATP-binding cassette (ABC) transporters. Nat Struct Mol Biol 23: 487–493. DOI: https://doi.org/10.1038/nsmb.3216, PMID: 27273632

Mi, W., Li, Y., Yoon, S.H., Ernst, R.K., Walz, T., Liao, M., 2017. Structural basis of MsbA-mediated lipopolysaccharide transport. Nature 549: 233–237. DOI: https://doi.org/10.1038/nature23649, PMID: 28869968

Oldham, M.L., Khare, D., Quiocho, F.A., Davidson, A.L., Chen, J., 2007. Crystal structure of a catalytic intermediate of the maltose transporter. Nature 450: 515–521.

Paoli, M., Marles-Wright, J., Smith, A., 2002. Structure-function relationships in heme-proteins. DNA Cell Biol 21: 271–280. DOI: https://doi.org/10.1089/104454902753759690

Perez, C., Gerber, S., Boilevin, J., Bucher, M., Darbre, T., Aebi, M., Reymond, J.L., Locher, K.P., 2015. Structure and mechanism of an active lipid-linked oligosaccharide flippase. Nature 524: 433–438. DOI: https://doi.org/10.1038/nature14953, PMID: 26266984

Pettersen, E.F., Goddard, T.D., Huang, C.C., Couch, G.S., Greenblatt, D.M., Meng, E.C., Ferrin, T.E., 2004. UCSF Chimera--a visualization system for exploratory research and analysis. J Comput Chem 25: 1605–1612. DOI: https://doi.org/10.1002/jcc.20084, PMID: 15264254

Procko, E., O’Mara, M.L., Bennett, W.F., Tieleman, D.P., Gaudet, R., 2009. The mechanism of ABC transporters: general lessons from structural and functional studies of an antigenic peptide transporter. FASEB J 23: 1287–1302. DOI: https://doi.org/10.1096/fj.08-121855, PMID: 19174475

Punjani, A., Rubinstein, J.L., Fleet, D.J., Brubaker, M.A., 2017. cryoSPARC: algorithms for rapid unsupervised cryo-EM structure determination. Nat Methods 14: 290–296. DOI: https://doi.org/10.1038/nmeth.4169, PMID: 28165473

Rempel, S., Gati, C., Nijland, M., Thangaratnarajah, C., Karyolaimos, A., de Gier, J.W., Guskov, A., Slotboom, D.J., 2020. A mycobacterial ABC transporter mediates the uptake of hydrophilic compounds. Nature 580: 409–412. DOI: https://doi.org/10.1038/s41586-0202072-8, PMID: 32296172

Rice, A.J., Park, A., Pinkett, H.W., 2014. Diversity in ABC transporters: type I, II and III importers. Crit Rev Biochem Mol Biol 49: 426–437. DOI: https://doi.org/10.3109/10409238.2014.953626, PMID: 25155087

Rodriguez, G.M., Smith, I., 2006. Identification of an ABC transporter required for iron acquisition and virulence in Mycobacterium tuberculosis. J Bacteriol 188: 424–430. DOI: https://doi.org/10.1128/JB.188.2.424-430.2006, PMID: 16385031

Ryndak, M.B., Wang, S., Smith, I., Rodriguez, G.M., 2010. The Mycobacterium tuberculosis high-affinity iron importer, IrtA, contains an FAD-binding domain. J Bacteriol 192: 861–869. DOI: https://doi.org/10.1128/JB.00223-09, PMID: 19948799

Skaar, E.P., 2010. The battle for iron between bacterial pathogens and their vertebrate hosts. PLoS Pathog 6: e1000949. DOI: https://doi.org/10.1371/journal.ppat.1000949, PMID: 20711357

Snow, G.A., White, A.J., 1969. Chemical and biological properties of mycobactins isolated from various mycobacteria. Biochem J 115: 1031–1050. DOI: https://doi.org/10.1042/bj1151031, PMID: 5360674

ter Beek, J., Guskov, A., Slotboom, D.J., 2014. Structural diversity of ABC transporters. J Gen Physiol 143: 419–435. DOI: https://doi.org/10.1085/jgp.201411164, PMID: 24638992

Wang, Z., Hu, W., Zheng, H., 2020. Pathogenic siderophore ABC importer YbtPQ adopts a surprising fold of exporter. Sci Adv 6: eaay7997. DOI: https://doi.org/10.1126/sciadv.aay7997, PMID: 32076651

Wells, R.M., Jones, C.M., Xi, Z., Speer, A., Danilchanka, O., Doornbos, K.S., Sun, P., Wu, F., Tian, C., Niederweis, M., 2013. Discovery of a siderophore export system essential for virulence of Mycobacterium tuberculosis. PLoS Pathog 9: e1003120. DOI: https://doi.org/10.1371/journal.ppat.1003120, PMID: 23431276

Xu, D., Feng, Z., Hou, W.T., Jiang, Y.L., Wang, L., Sun, L., Zhou, C.Z., Chen, Y., 2019. Cryo-EM structure of human lysosomal cobalamin exporter ABCD4. Cell Res 29: 1039–1041. DOI: https://doi.org/10.1038/s41422-019-0222-z, PMID: 31467407

Yang, R., Cui, L., Hou, Y.X., Riordan, J.R., Chang, X.B., 2003. ATP binding to the first nucleotide binding domain of multidrug resistance-associated protein plays a regulatory role at low nucleotide concentration, whereas ATP hydrolysis at the second plays a dominant role in ATP-dependent leukotriene C4 transport. J Biol Chem 278: 30764–30771. DOI: https://doi.org/10.1074/jbc.M304118200, PMID: 12783859

Yang, X., Hu, T., Yang, X., Xu, W., Yang, H., Guddat, L.W., Zhang, B., Rao, Z., 2020. Structural Basis for the Inhibition of Mycobacterial MmpL3 by NITD-349 and SPIRO. J Mol Biol 432: 4426–4434. DOI: https://doi.org/10.1016/j.jmb.2020.05.019, PMID: 32512002

Zhang, B., Li, J., Yang, X., Wu, L., Zhang, J., Yang, Y., Zhao, Y., Zhang, L., Yang, X., Yang, X., Cheng, X., Liu, Z., Jiang, B., Jiang, H., Guddat, L.W., Yang, H., Rao, Z., 2019. Crystal Structures of Membrane Transporter MmpL3, an Anti-TB Drug Target. Cell 176: 636–648 e613. DOI: https://doi.org/10.1016/j.cell.2019.01.003, PMID: 30682372

Zheng, S.Q., Palovcak, E., Armache, J.P., Verba, K.A., Cheng, Y., Agard, D.A., 2017. MotionCor2: anisotropic correction of beam-induced motion for improved cryo-electron microscopy. Nat Methods 14: 331–332. DOI: https://doi.org/10.1038/nmeth.4193, PMID: 28250466

